# Simulated brain networks reflecting progression of Parkinson’s disease

**DOI:** 10.1101/2024.01.12.574450

**Authors:** Kyesam Jung, Simon B. Eickhoff, Julian Caspers, UKD-PD team, Oleksandr V. Popovych

## Abstract

Neurodegenerative progression of Parkinson’s disease affects brain structure and function and, concomitantly, alters topological properties of brain networks. The network alteration accompanied with motor impairment and duration of the disease is not yet clearly demonstrated in the disease progression. In this study, we aim at resolving this problem with a modeling approach based on large-scale brain networks from cross-sectional MRI data. Optimizing whole-brain simulation models allows us to discover brain networks showing unexplored relationships with clinical variables. We observe that simulated brain networks exhibit significant differences between healthy controls (*n*=51) and patients with Parkinson’s disease (*n*=60) and strongly correlate with disease severity and disease duration of the patients. Moreover, the modeling results outperform the empirical brain networks in these clinical measures. Consequently, this study demonstrates that utilizing simulated brain networks provides an enhanced view on network alterations in the progression of motor impairment and potential biomarkers for clinical indices.

## Introduction

Parkinson’s disease (PD) is associated with degeneration of dopaminergic neurons in the substantia nigra pars compacta^1^. This dopamine deficiency involved in basal ganglia circuits leads to movement disorder^2^. As a neurodegenerative disease, PD progresses over time. Evidently, the disease duration is associated with severity of motor impairment^3^. Accordingly, taking care of the symptom severity after the disease onset is crucial to quality of patients’ life. Medication with levodopa or dopaminergic therapy is an effective treatment of PD without diminished effects in a long disease duration of decades^4,5.^ However, with such a long-time neurologic state before and after the diagnosis, it may cause corresponding irreversible brain network alteration ^6,7^ as well as drug-induced dyskinesia^8^ or cognitive deficits^9^. These degenerative alterations are important for understanding the progression of the disease in pre-motor or prodromal periods (before the disease onset), which is one of clinical challenges.

Investigating the changes of brain networks with the disease development can help to understand its pathological progress. Eventually, these changes do not only impact whole-brain networks, but also correspondingly alter their topological properties of the networks. Network representation of the human brain, also known as human connectome^10^, has widely been employed in neuroscientific research for understanding the neural coding and information transmission via brain circuits^11^. Human connectome comprises various network models^12^, and graph theory provides effective tools to evaluate properties of such complex networks from large-scale brain connectivity^13^. Many studies have addressed the relationships between network properties and behavior^14^ including diseased states^15-17^. In other words, the network-based approach provides features reflecting psychological and clinical attributes^18^. In particular, brain networks of PD patients have been shown to be different from those of healthy participants in network integration and segregation^19,20.^ Besides, it can also be employed for patient classification^21-23^.

Examining the network alterations through experimental interventions for *in vivo* human brain is hardly feasible. On the other hand, *in silico* brain networks derived by whole-brain modeling have no limitation in virtual interventions for instance virtual corpus callosotomy^24^. In this study we therefore suggest an approach that utilizes whole-brain dynamical models and resultant simulated brain networks for enhanced relationship of them with clinical variables. With this, we probe simulated network properties by varying model parameters and search for optimal values that provide the best simulation model correspondence to research objectives^22^. This behavioral network-based model fitting is a novel approach to investigate the relationships between simulated network properties and clinical measures. It allows us to explore *in silico* brain dynamics for study conditions that are not available for the analysis of empirical human data *in vivo*.

Here we test the relationship between simulated brain networks and clinical scores related with the progression of PD, *i.e*., disease severity and disease duration. For the severity, we utilize the unified Parkinson’s disease rating scale (UPDRS)^25^ in the modeling approach. It is also used to infer the effect of medication (dopaminergic therapy) on motor impairment with the extent of striatal dopamine depletion^7^. Accordingly, we demonstrate an important and intriguing dependence of the modeling results on clinical variables by applying the behavioral network-based model fitting. We in particular opt for network modularity (segregation) and efficiency (integration) and address the relationship between the network properties and clinical variables of PD (Fig. 1). As a result, we demonstrate significant differences of the simulated network properties between PD patients and healthy controls as well as correlations with disease severity and duration, which were found to be unclear in empirical brain networks. Our results therefore reveal that simulated brain networks clearly reflect the clinical properties of the disease, and the suggested model-fitting approach contributes to investigation and better understanding the disease progression. In consequence, it will possibly provide a new observer-independent biomarker for disease progression.

**Figure 1.**
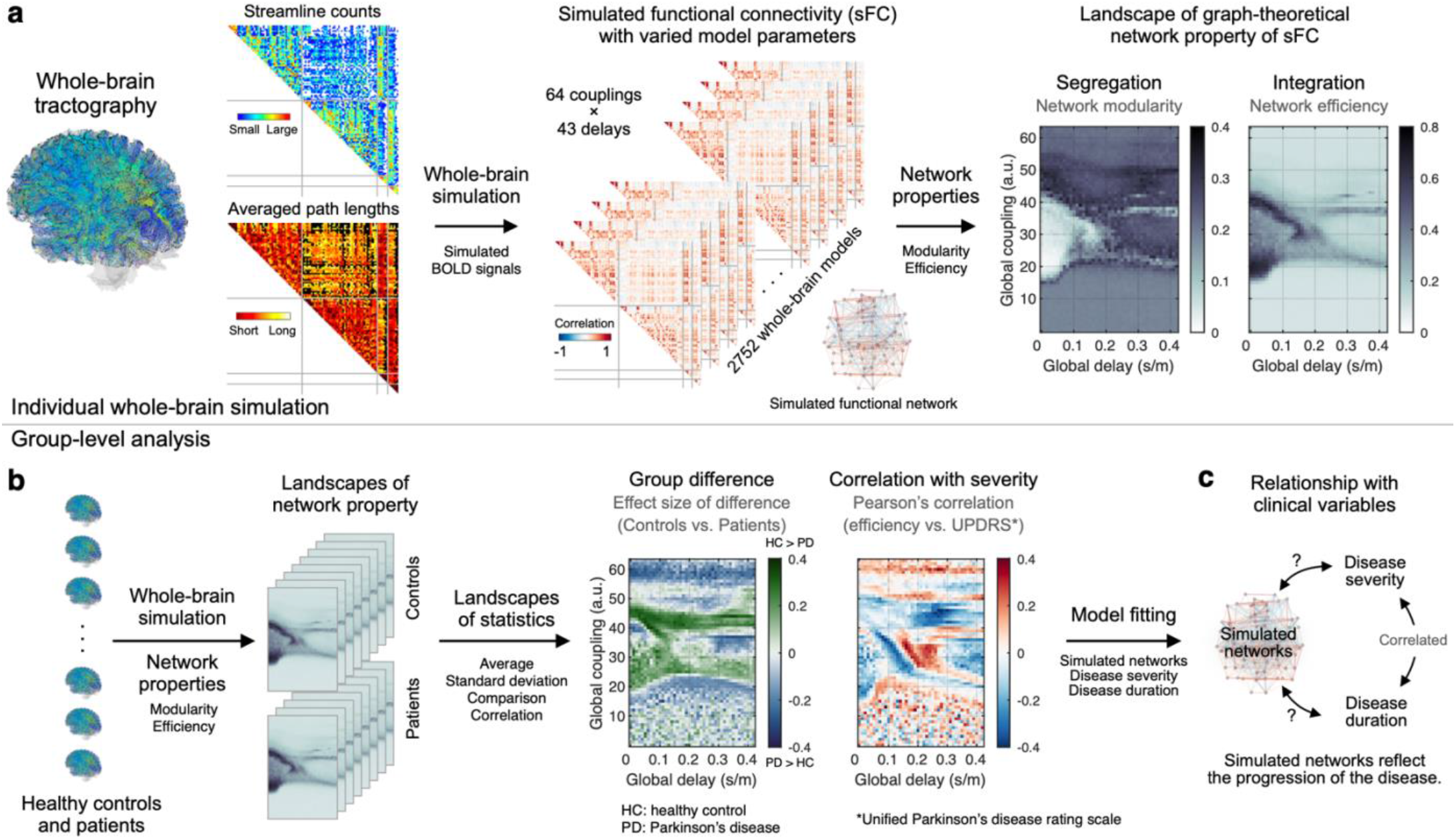
Workflow of the study. **(a)** Individual whole-brain tractography calculated from diffusion-weighted MRI data was used to extract parcellation-based empirical structural connectivity (streamline counts and streamline path lengths) that was then utilized for derivation of the whole-brain dynamical model, simulation of the resting-state brain activity and calculation of simulated functional connectivity for varying model parameters. For every subject a few parameter landscapes were obtained representing the properties of the simulated FC networks, *e.g*., network modularity and efficiency versus model parameters. **(b)** Individual parameter landscapes of network properties were used for group-level analysis to obtain parameter landscape of statistics of simulated network properties across subjects, *e.g*., group differences between patients and healthy controls and correlation between disease severity and network properties. **(c)** The statistical parameter landscapes and results of the behavioral model fitting were employed for investigation of the relationships between the simulated network properties and clinical variables, *e.g*., disease severity and duration.

## Results

### Relationships between clinical variables

We calculated Pearson’s correlation coefficients between clinical and demographic variables. Severity of motor impairment based on the UPDRS III scores does not significantly correlate with age, disease onset age, and disease duration in the considered cross-sectional data (Table 1). We observed a weak positive correlation between UPDRS III Off (condition without medication) and disease duration. This tendency is consistent with an increase of UPDRS III Off with disease duration in a longitudinal study^3^. Instead, UPDRS III Off and UPDRS III On (condition with medication) strongly correlate with each other. As for the effect of medication, the within-subject difference of UPDRS III (Off – On) shows significant correlations with the onset age and the duration of the disease, but it is not the case for age. For a progressive disease, the disease duration is obviously a more important factor to delve into the effect of medication compared to age.

**Table 1.**
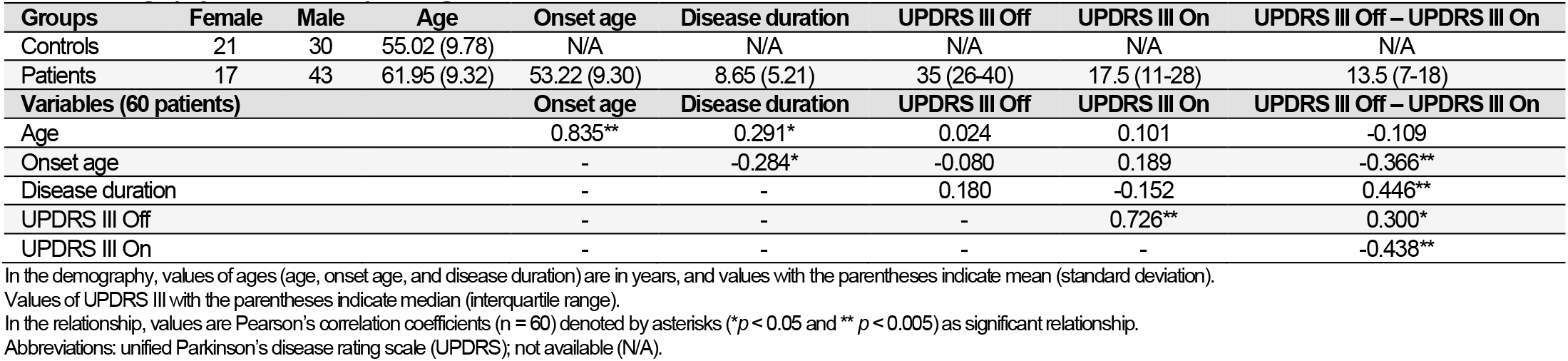
Demography and relationship among clinical variables.

### Whole-brain model fitting using network landscapes

We calculated network modularity and efficiency of the simulated functional connectivity (FC) and projected them on a parameter space that comprises model parameters of global couplings and delays (Fig. 1a). Each subject therefore has landscapes of network modularity and efficiency. By the group-level (across subjects) statistical analyses (Fig. 1b) we obtained the respective statistical parameter maps (Fig. 2). The latter in particular include parameter regimes showing significant results and relatively large inter-subject variability of network properties for robust performance against noise (Suppl. Fig. 1a-b). The intersections of these regimes were used for the behavioral network-based model fitting as a parameter mask (Suppl. Fig. 1c). Then, we searched for the optimal parameter points (the magenta-white squares in Fig. 2) corresponding to the largest effect size of group difference between healthy controls and patients (Fig. 2a-b) and the strongest positive or negative Pearson’s correlation coefficients between the network properties and UPDRS III On scores (Fig. 2c-e). The optimal global delays were located in the interval [0.06, 0.25] s/m of biologically feasible signal propagation delays^26^ (squares between vertical lines in Fig. 2). We further clarified reliability of the model fitting via cross-validated model fitting^22^. As a result, the selected parameter points (squares in Fig. 2c-e) were stable across different subject configurations by random sampling (Suppl. Fig. 2).

**Figure 2.**
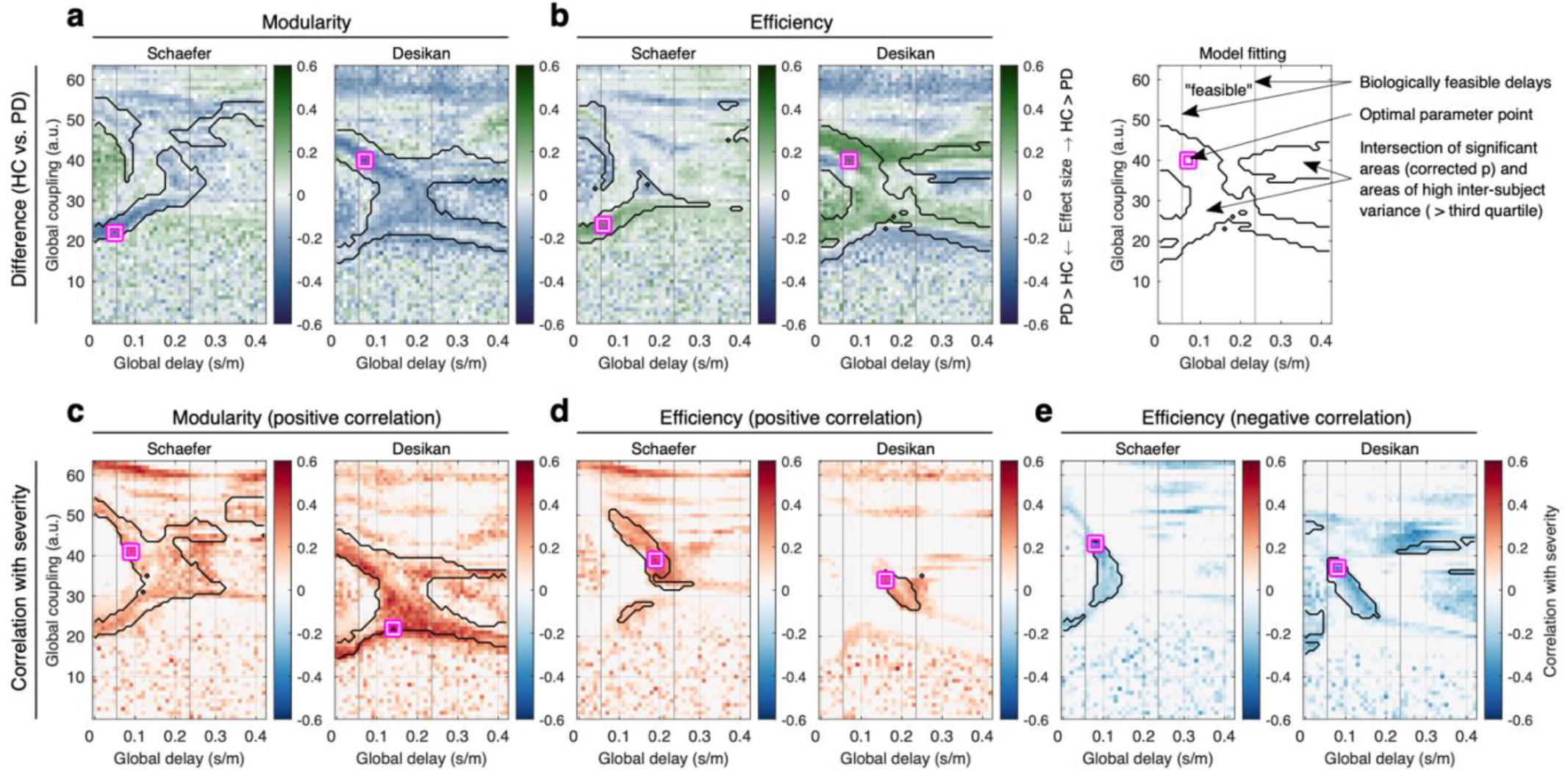
Parameter landscapes of behavioral model fitting of the whole-brain dynamical model of the Jansen-Rit type. The network modularity (functional segregation) and network efficiency (functional integration) of simulated FC were used to calculate **(a-b)** landscapes of the effect size of the group difference between healthy controls (HC) and patients with Parkinson’s disease (PD) and **(c-e)** landscapes of Pearson’s correlation (across PD patients) between simulated network properties and severity of the disease as given by the unified PD rating scales (UPDRS III medication On). The calculations were performed for the Schaefer and the Desikan-Killiany (Desikan) brain atlases indicated in the titles of plots together with the respective network properties. The color depicts the effect size and correlation in plots (a-b) and (c-e), respectively. The vertical lines bound an approximate range of biologically feasible delays, the magenta-white squares indicate optimal parameter points of the largest effect size or correlation in the parameter domain bounded by the black contour curves of intersection between significant areas thresholded by the random-field theory for multiple tests and areas of high inter-subject variance of the respective network properties (> third quartile), see the rightmost plot in the upper row for explanation.

Landscape of the group difference shows that network modularity of PD patients is mostly higher than that of controls (Fig. 2a). In contrast, network efficiency of the patients is lower than that of the controls (Fig. 2b). The landscape of the correlations between the network modularity and the disease severity mostly shows positive correlations (Fig. 2c). Remarkably, the network efficiency of simulated FC alters a lot by varying the model parameters on the landscape leading to positive and negative correlations (Fig. 2d-e). With these landscape patterns, switching the optimal parameters from small to large delays impacts brain dynamics leading to opposite tendencies of correlations. Therefore, we inferred that simulated functional networks of patients may comparatively change when model parameters are varied, and simultaneously, the changes of the network property can be related with the severity of the disease.

### Group difference of network properties

We also calculated network properties of the empirical connectomes, *i.e*., FC and structural connectivity (SC), and compared the network properties of healthy controls with those of patients. We find no significant group differences in the empirical data (Fig. 3a-b). In contrast, the network properties of the simulated FCs obtained for the optimal model parameters (squares in Fig. 2a-b) show significantly different distributions between the controls and the patients. The modularity of patients is higher, and the efficiency is lower than those of healthy controls for both parcellations (Fig. 3c), which is consistent with the tendency reported in the literature for empirical neuroimaging data^20^. Based on these findings, we conclude that the modeling results are different for healthy controls and PD patients. As a consequence, segregation (modularity) and integration (efficiency) of simulated functional networks can be engaged in the relationship with clinical variables of PD, which we consider in detail below.

**Figure 3.**
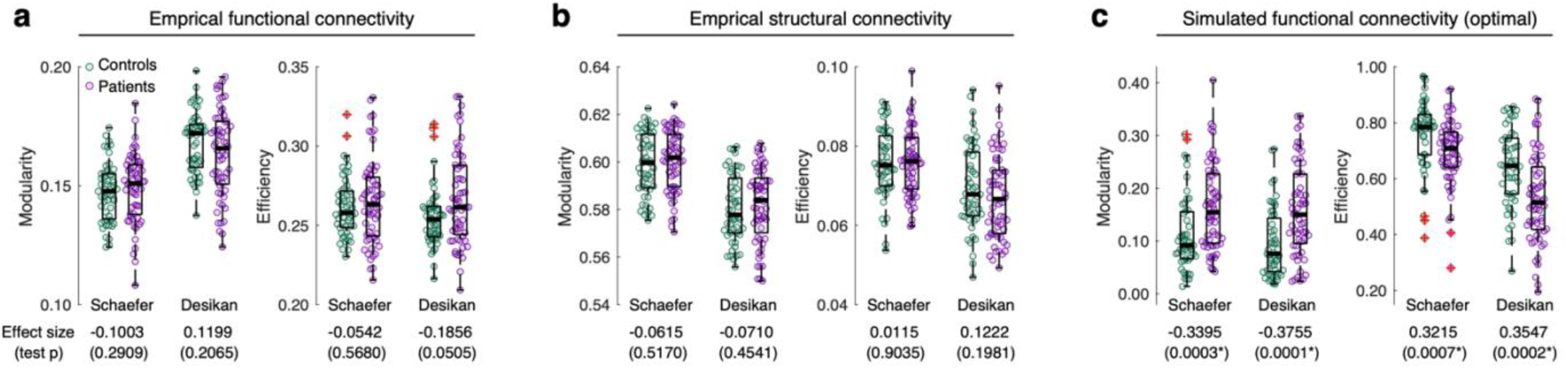
Group difference between healthy controls (HC, *n*=51) and patients with Parkinson’s disease (PD, *n*=60) for two brain parcellations and several comparison conditions. **(a-c)** The group differences of the network modularity and efficiency between HC and PD for **(a)** empirical functional connectivity, **(b)** empirical structural connectivity, and **(c)** optimal simulated functional connectivity. The empty circles in the plots correspond to individual subjects. The brain parcellations are indicated in the plots and the values under the plots are the effect sizes of the group difference (positive for HC > PD and negative for PD > HC) and their statistics (*p*-values of the Wilcoxon rank-sum two-tail test). The *p-*values with asterisks indicate significant results (*p* < 0.05). The middle thick lines in the interquartile boxes indicate the medians of distributions, and the red crosses are outliers.

### Simulated network property reflects severity of the disease

We calculated Pearson’s correlation coefficients between the network properties (modularity and efficiency) and the disease severity (UPDRS III On). The network properties of empirical FC and SC do not show significant correlation with disease severity (Fig. 4a-d, Table 2). On the other hand, the network properties of simulated FC obtained from the model simulations with the optimal parameters (squares in Fig. 2c-e) significantly correlate with the disease severity (Fig. 4e-g). Interestingly, the simulated network efficiency clearly shows opposite tendencies, *i.e*., positive and negative correlations with the disease severity (Fig. 4f-g). In the network topology of whole-brain connectivity^13^, high segregation (modularity) corresponds to the presence of segregated (weakly interacting) network modules of densely interconnected nodes (brain regions) within modules. On the other hand, high integration (efficiency) is related with strong connections also between modules. Therefore, with the above results, we arrived at an assertion that the network-based modeling discloses the severity of motor impairment of PD patients via probing into the segregation and integration of simulated brain networks.

**Table 2.** Correlation between empirical network properties and clinical variables (60 patients).

| Empirical functional connectivity | Age | Onset age | Disease duration | UPDRS III Off | UPDRS III On | UPDRS III Off – UPDRS III On |
| --- | --- | --- | --- | --- | --- | --- |
| Modularity (Schaefer) | -0.047 | -0.143 | 0.166 | -0.144 | -0.172 | 0.050 |
| Modularity (Desikan-Killiany) | -0.346* | -0.293* | -0.094 | -0.072 | -0.133 | 0.090 |
| Efficiency (Schaefer) | 0.287* | 0.332* | -0.078 | -0.023 | 0.095 | -0.162 |
| Efficiency (Desikan-Killiany) | 0.358** | 0.377** | -0.032 | -0.014 | 0.080 | -0.129 |
| Empirical structural connectivity | Age | Onset age | Disease duration | UPDRS III Off | UPDRS III On | UPDRS III Off – UPDRS III On |
| Modularity (Schaefer) | 0.230* | 0.063 | 0.291* | 0.159 | 0.183 | -0.046 |
| Modularity (Desikan-Killiany) | 0.246 | 0.068 | 0.310* | 0.170 | 0.186 | -0.036 |
| Efficiency (Schaefer) | -0.209 | -0.054 | -0.270* | -0.219 | -0.204 | -0.003 |
| Efficiency (Desikan-Killiany) | -0.073 | 0.018 | -0.158 | -0.206 | -0.149 | -0.064 |

In the relationship, values are Pearson's correlation coefficients ( $n = 60$ ) denoted by asterisks (\* $p < 0.05$ and \*\* $p < 0.005$ ) as significant relationship. Abbreviations: unified Parkinson's disease rating scale (UPDRS).

**Figure 4.**
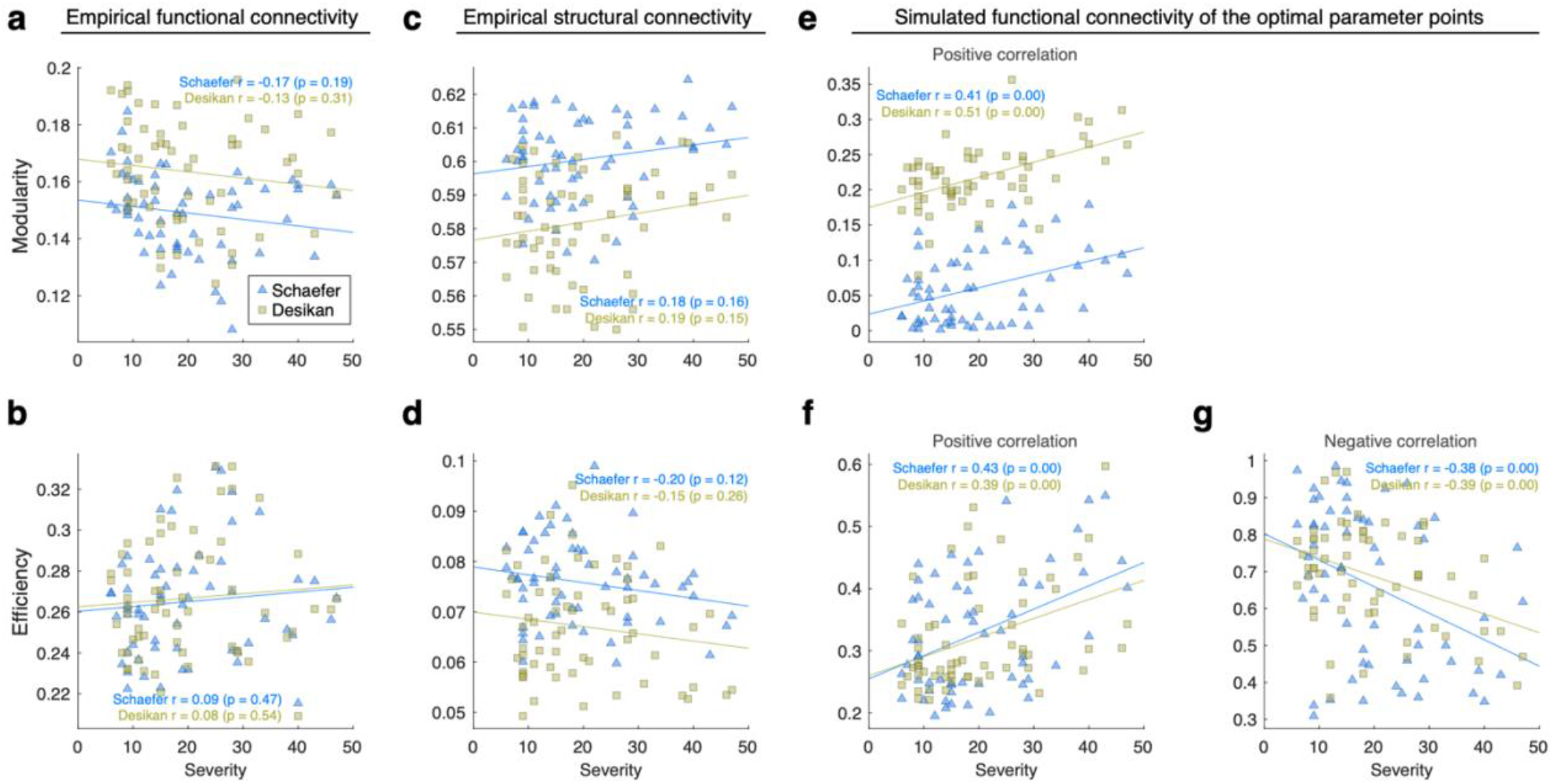
Correlation between severity of the disease of individual PD patients as given by the unified Parkinson’s disease rating scales (UPDRS III medication On, horizontal axes) and modularity and efficiency network properties (vertical axes) of empirical and simulated brain connectomes for **(a-b)** empirical FC, **(c-d)** empirical SC and **(e-g)** simulated FC calculated for the optimal parameters denoted by the squares in Fig. 2c-e. The depicting triangles and squares in the plots denote the two considered brain parcellations correspond to individual PD patients. The used brain parcellations, calculated Pearson’s correlation coefficient and its statistical test (*p*-value) are indicated in the legends.

To investigate the changes of the simulated brain network topology, we in detail analyzed connections (network edges) of simulated FC across the patients. To do this, we considered the optimal models corresponding to the strongest negative and positive correlations of the network efficiency with the disease severity (squares in Fig. 2d-e). In the case of small delay (negative correlation), many inter-hemispheric connections significantly correlate with the disease severity as compared with the intra-hemispheric connections (lower triangles in Fig. 5a-b). On the other hand, in the case of large delay (positive correlation), many intra-hemispheric connections significantly correlate with the disease severity (upper triangles in Fig. 5a-b). Based on this result, we opted for the significant edges of the simulated FC from the small delays as edges of interest (non-white edges in lower triangles in Fig. 5a-b) for further analysis. Thereafter, we subtracted the selected FC edges of small delay from the corresponding FC edges of large delay for every subject and related these edge differences to the disease severity of a given subject. As a result, we can clearly see that the individual histograms of the edge differences are away from zero when the patients have less severe motor impairment and approach the origin when the severity increases (Fig. 5c-d). Accordingly, the medians of the edge differences exhibit the same behavior and significantly correlate with the disease severity (Fig. 5e). Consequently, changes of the corresponding network efficiency also significantly correlate with the disease severity (Fig. 5f). Furthermore, the network efficiency of healthy controls significantly and strongly deviates from zero as a possible limiting case of zero disease severity (left most boxplots in Fig. 5f). Thus, our results attest that simulated brain networks of patients with severe symptoms show less changes in the network integration respecting to the network alterations of the models with distinct optimal parameter points.

**Figure 5.**
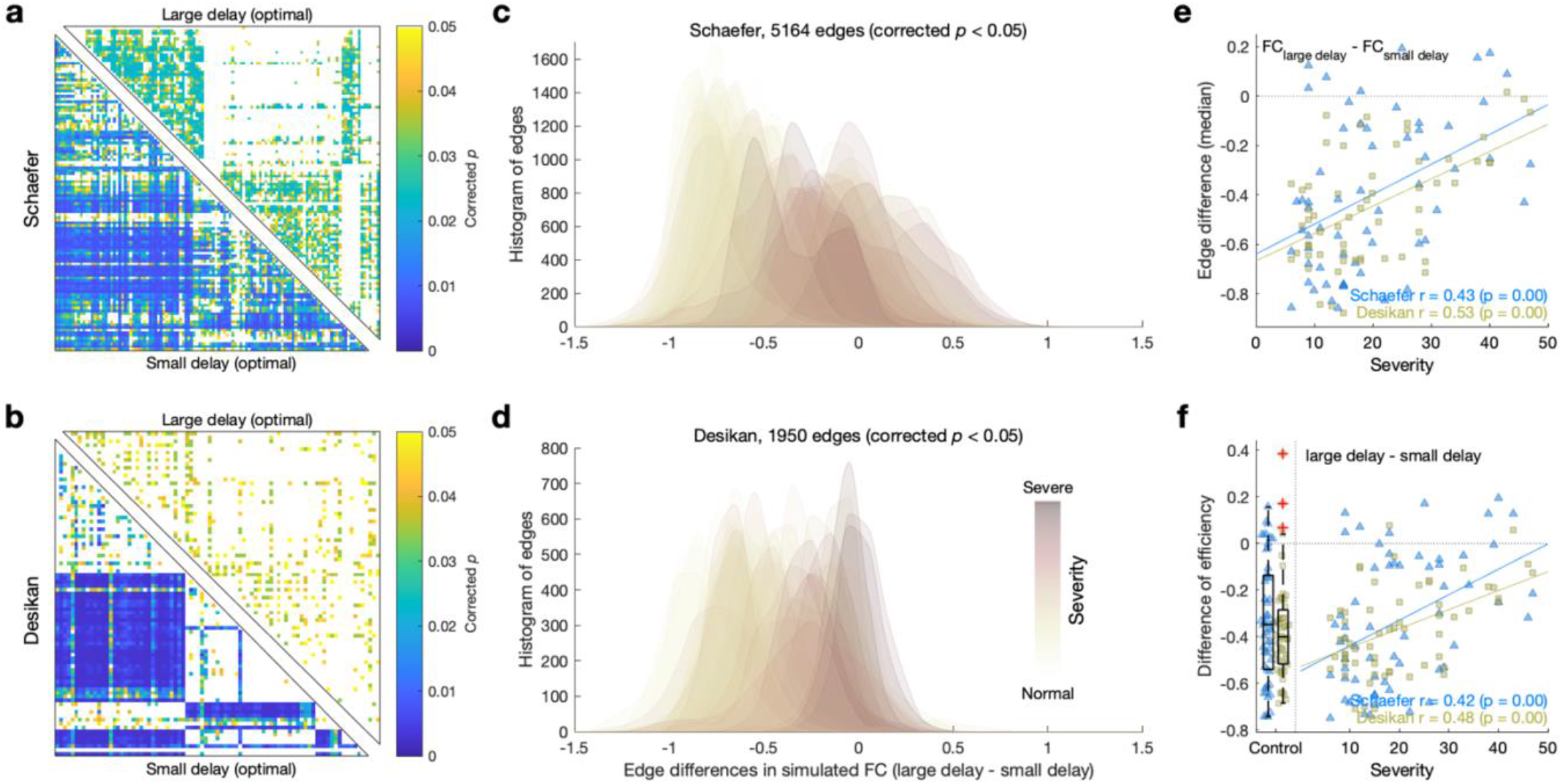
Correlations between disease severity as given by the unified Parkinson’s disease rating scales (UPDRS III, medication On) and simulated functional connectivity (FC). The latter was simulated for the optimal model parameters of the strongest positive and negative correlations between the disease severity and network efficiency of simulated FC obtained for large and small optimal delays, respectively (Fig. 2d-e, square marks). **(a-b)** Results of statistical tests (*p*-values corrected by the Benjamini-Hochberg false discovery rate) of Pearson’s correlation (across patients) between the disease severity and the edges of the simulated FC for the optimal model parameters of small (lower triangles) and large (upper triangles) delays for each parcellation indicated at the left. **(c-d)** Histograms of the differences of significant FC edges from lower triangles of the corrected *p* matrices (a-b) between the large and small optimal delays of each patient and parcellation scheme. The color shading of the histograms indicates the severity of the disease of corresponding patients. **(e-f)** Scatter plots of the relationships between the disease severity and the differences between the values for the large and small delays of **(e)** the medians of the histograms in (c-d) and **(f)** the network efficiency. The depicting triangles and squares in the plots denote the two considered brain parcellations correspond to individual PD patients. The amount of correlation of the depicted relationships are indicated in the plots together with the results of its statistical tests (*p*-values) of Pearson’s correlation for both considered parcellation schemes. In the scatter plot (f, left side), the box plots depict the distributions of the respective values of the efficiency differences (vertical axes) for 51 healthy controls, where the middle lines in the interquartile boxes indicate the medians of distributions, and the red crosses are outliers. Both distributions were normally distributed according to the Kolmogorov-Smirnov normal distribution test and significantly different from zero (one-sample two-tail *t*-test).

### Effect of medication and network efficiency relative to disease duration

The severity of motor impairment varies across patients in the considered cohort, and UPDRS III Off scores are higher than UPDRS III On scores (Suppl. Fig. 3a). The difference between the scores (Off – On) can be considered as an effect of medication on the symptoms, which exhibits significant correlation with the disease duration (Table 1, Suppl. Fig. 3b).

Empirical FC network properties do not show significant correlation with the disease duration, but those of empirical SC network are significantly correlated with disease duration (Table 2). To evaluate the impact of the disease duration on the simulated FC network, we applied the behavioral network-based model fitting to the disease duration instead of the disease severity (Suppl. Fig. 4a-b). Network modularity of simulated FC for the optimal parameter points demonstrates positive correlations with the disease duration, which is consistent with the case of empirical SC network (Table 2).

Interestingly, we found parameter regimes of negative and positive correlations of the network properties with the disease duration (Fig. 6a), where the landscapes of the correlations of the network efficiency with the disease severity show similar patterns with each other (Suppl. Fig. 4b, compare to Fig. 2d-e). This is in spite of a weak negative correlation between the disease duration and severity (UPDRS III On) in clinical measures (Table 1). For the alterations of network efficiency between negative and positive correlations (Fig. 6b-c), we calculated a ratio of the network efficiency of large delay to that of small delay and related it to the disease duration (Fig. 6d). The network efficiency appeared to be smaller for the case of large delay when the patient has a relatively short disease duration. Accordingly, the ratio, an alteration of network efficiency, significantly correlates with disease duration (Fig. 6d-e).

**Figure 6.**
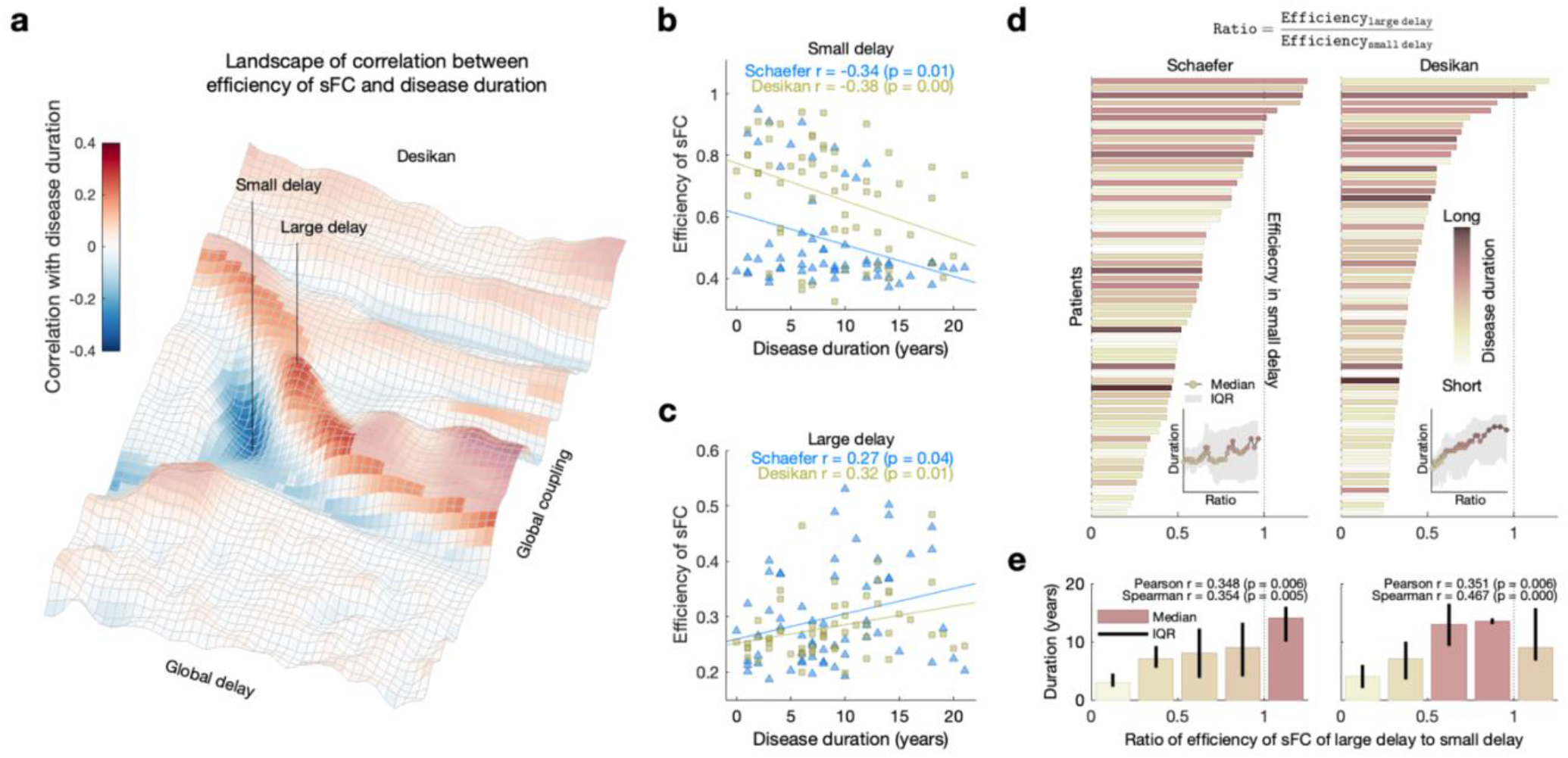
Relationships between network efficiency of simulated FC and disease duration. **(a)** Parameter landscape of Pearson’s correlation coefficients between simulated network efficiency and the disease duration in the Desikan-Killiany (Desikan) atlas. The vertical lines with ‘small delay’ and ‘large delay’ indicate optimal parameter points for negative and positive correlation, respectively. **(b-c)** Scatter plots for **(b)** negative and **(c)** positive correlations between disease duration and network efficiency of the simulated FC at the optimal parameter points with small and large delay, respectively. The lines are linear fitting between simulated network efficiency and the disease duration. The depicting triangles and squares in the plots denote the two considered brain parcellations correspond to individual PD patients. The amount of correlation of the depicted relationships are indicated in the plots together with results of statistical tests (*p*-values) for both considered parcellation schemes. **(d)** Ratio of the optimal simulated network efficiency of large delay to that of small delay for individual patients (vertical axes) sorted according to the ratio of the network efficiency indicated by the horizontal bars with color depicting the disease durations of the corresponding patients. The inserts show medians of the disease duration corresponding to the moving average along the patients in ascending order of the efficiency ratio. The gray shadow indicates inter-quartile ranges (IQR) of the disease duration. **(e)** Bar plots of the median values of the efficiency ratio with error bars indicating IQR of ratios in five intervals splitting the range from 0 to 1.25. The amount of correlation of the depicted relationships between the efficiency ratio and the disease duration are denoted in the plots together with results of statistical tests (*p*-values) of the Pearson’s correlation and the Spearman’s correlation, respectively.

## Discussion

The aim of the current study was to demonstrate the relationship between simulated brain network properties and clinical variables considering the progression of PD, such as severity of motor impairment and disease duration. The reported results indicate that functional segregation and integration of simulated brain networks can significantly correlate with disease severity (UPDRS III). In addition, the simulated network properties provide higher effect sizes of the group difference between healthy controls and PD patients than those of empirical networks. Remarkably, alterations of efficiencies of simulated brain networks derived by the models with distinct optimal parameters evidently reflect the clinical measures (the severity and duration of the disease). As a potential approach, the behavioral network-based model fitting allows us to explore simulated brain networks that reflect the progression of the disease. Consequently, we suggest a way how to generate and utilize simulated human connectomes for investigation of behavioral or clinical measures as well as disease onset and progression.

The results of the current study indicate that simulated data of whole-brain dynamical models can provide features of brain networks with a great potential for enhancing group differences between healthy controls and patients with PD. Furthermore, the simulated brain networks clearly reflect clinical scores such as disease severity and duration. Based on the modeling results, the simulated brain networks outperform the empirical networks in regard to group differences and correlations with clinical variables. In contrast, empirical FC did not exhibit such clear relationships of the modeling, and empirical SC only showed significant correlation with disease duration (Table 2). Thus, the whole-brain modeling is essential to reveal relationships between brain networks and clinical measures. In contrast to using empirical brain networks, the data-driven modeling approach^27^ allows us to explore model parameters and search for the most effective models that simulate brain dynamics at the best correspondence to the posed research questions.

The simulation results for the functional integration (network efficiency) are more involved with the disease severity than that of the functional segregation (network modularity) because the network efficiency does not only show negative correlations (Fig. 4f) but also have positive correlations (Fig. 4g) with disease severity. In addition, it exhibits regimes that are associated with alteration of the inter-hemispheric connections in simulated brain networks. In other words, inter-hemispheric connections in the brain of the patients with severe motor impairment are not adequately contributing to whole-brain dynamics. This interpretation refers to the literature addressed that the inter-hemispheric connections are related with the functional integration^28^. Besides, decreased inter-hemispheric connections in PD were also observed in the empirical SC in the current study as well (Suppl. Fig. 5). This is consistent with the empirical result of decreased resting-state inter-hemispheric connections in PD^29^.

The severity of motor impairment of PD is expected to increase over time^3^ because of the progressive disease. Although we observe a week correlation between the severity and duration of the disease in these cross-sectional clinical scores (Table 1), instead we interpret our last modeling results (Figs. 5 and 6) as the impact of the disease progression on the simulated network efficiency, which reflects the disease severity and duration. In addition, the positive correlation between the difference of UPDRS III scores (Off – On) and the disease duration (Suppl. Fig. 3b) allows us to interpret the effect of medication as being related with the disease duration^9^. With this, we infer that the alteration of the simulated network efficiency can intermediate the relationship of the disease duration with the effect of medication. Therefore, the simulation outcomes of the dynamical models can be used to integrate the relationships among the disease progression (disease severity and duration) and the effect of medication via interrelating them with topological properties of simulated brain networks (see Fig. 1c). In summary, our results support the claim that the whole-brain dynamical modeling can provide a potential way for understanding the interrelations between the properties of brain networks and PD progression and severity.

One of the main advances of the current novel approach is that the calculated parameter landscapes of simulated network properties provide statistical maps and bases for the behavioral model fitting. Subsequently, significant regimes in a landscape can be selected for the best correspondence to target variables as for research questions. This analysis has some analogy with statistical brain mapping in neuroimaging research related with behavioral tasks^30^, clinical tests^31^, genetic measures^32^, and so forth. Therefore, we can also apply this approach to the parameter landscapes (mapping) of the simulated brain dynamics that can be related with behavioral and clinical measures as we demonstrated in this study.

The modeling and validation methods employed in this study rely on empirical SC and clinical variables, instead of empirical FC as a target variable. This may have a few advantages including a possible positive influence on reliability of simulation results. Indeed, empirical FC and SC have different test-retest reliabilities, where the empirical resting-state FC showed reliability in the range from 0.3 to 0.6 (multi-sites with various scan times)^33-35^ in intraclass correlation coefficient (ICC). On the other hand, ICC values of empirical SC were found between 0.7 and 0.8 (intra-site, inter-site, and multi-sites)^36,37.^ Accordingly, network properties of empirical SC also showed a higher reliability than that of empirical resting-state FC^38^. With such different reliabilities, the current modeling approach could also show varied results when we use different study conditions. Therefore, we applied a cross-validated model fitting approach^22^ to the landscapes of correlations between simulated network efficiency and disease severity. As a result, it shows stable optimal model parameter points across different patient configurations by random sampling (Suppl. Fig. 2). Therefore, the suggested behavioral network-based model fitting may show consistent outcomes when we include unseen subjects into the analysis.

The current whole-brain model considers the excitatory and inhibitory populations in each brain region and (local and global) interactions among them which represents neurophysiological activities in coupled cortical columns^39^. However, the current model is still relatively simple compared to the network from such a complex human brain. Thus, it is necessary to investigate individualized whole-brain models with varied local parameters instead of fixed values such as excitatory-inhibitory balances in the future study. For generalized modeling, the current approach needs to be evaluated by including multi-site data.

Investigating alterations of the brain connectome is essential for understanding progression of neurodegenerative diseases. Respectively, our findings can be utilized for future research that can show the impact of the disease duration on the whole-brain dynamics. Furthermore, including prodromal subjects and longitudinal data will provide a way of the validation of the current approach as progressive markers. Therefore, future investigations about the impact of the progression of PD on disease symptoms and brain networks will use longitudinal data that consists of healthy, prodromal, and diseased subjects from multi-clinical sites. For an advanced approach, we can consider individualized whole-brain models with varied local parameters using high-dimensional parameter optimization^40^ and its applications, for instance, a computational biomarker for individual clinical scores. As a consequence, the modeling outcome can be used for objective evaluations of these clinical indices.

## Methods

### Participants

Multimodal brain MRI data were acquired in 111 human subjects including 51 healthy subjects (21 females; age range: 41-78 years) and 60 patients (17 females; age range: 44-80 years) with Parkinson’s disease (PD). Age of symptom onset as well as disease duration were acquired in PD patients. Furthermore, the unified Parkinson’s disease rating scales (UPDRS)^25^ was assessed by an expert neurologist for each patient in a condition under regular medication (UPDRS-On) as well as after at least 12 hours withdrawal of all dopaminergic drugs (UPDRS-Off). All healthy controls had no history of any neurologic or psychiatric disease and no abnormalities of cranial MRI. The study was approved by the local ethics committee and performed in accordance with the Declaration of Helsinki. Written informed consent was obtained prior to study inclusion.

### MRI protocols and processing

A 3T scanner (Siemens Trio) was used for T1-weighted MRI (T1w; voxel size = 1.0×1.0×1.1 mm^3^), diffusion-weighted MRI (dwMRI; B = 1000 s/mm^2^ with 64 directions; voxel size = 2.4×2.4×2.4 mm^3^) with a non-weighted (B0) image, and resting-state functional MRI (rs-fMRI; repetition time = 2.21 s; 300 volumes during 663 s; voxel size = 3.125×3.125×3.565 mm^3^). MRI processing was performed using a pipeline that included structural and functional modules. The structural module performed preprocessing for T1w (bias-field correction; alignment of anterior-posterior commissures; brain tissue segmentation; reconstruction of gray-white matter boundary and pial surface) and dwMRI (removing the Gibbs ringing effect; correction of bias-field, head-motion, and eddy distortion). Subsequently, whole-brain tractography (WBT) with 10 million streamlines was calculated based on estimated fiber orientation distributions of white matter using spherical deconvolution^41,42.^ The functional module processed rs-fMRI (correction of slice-time and head-motion; re-slicing in a 2 mm iso-cubic voxel space; intensity normalization; de-trending; nuisance regression with regressors of white matter, cerebrospinal fluid, the entire brain, and the head motion). Two brain atlases were used for cortical parcellation based on the Schaefer^43^ atlas with 100 parcels and Desikan-Killiany^44^ atlas with 68 parcels.

Subcortical regions were also added into each atlas. The pipeline counted streamlines in WBT connecting any two brain regions, which shaped SC. It also extracted mean blood oxygenation level-dependent (BOLD) signals of each brain region from the processed rs-fMRI and calculated the Pearson’s correlation coefficients between the BOLD signals of any two brain regions, which constitutes FC. More details of the MRI processing and tracking parameters can be found elsewhere^22^.

### Empirical network properties

For the empirical functional network properties, we took absolute values of edges in empirical FC and used the brain connectivity toolbox to calculate network modularity and global efficiency^13^. For the empirical structural network properties, streamline counts of edges in empirical SC were divided by the maximal number of streamlines (self-connections were excluded) and used for the network properties as well.

### Neural mass model of two neural populations

Post-synaptic potentials (PSP) were simulated via interacting between excitatory and inhibitory neural populations in each brain region based on Jansen-Rit model type^39,45^ as a convolution-based neural mass model^46^ transforming the average of pre-synaptic firing density into the average PSP between excitatory neural populations in brain regions.

Simulated PSP signals were obtained by using the following differential equations:

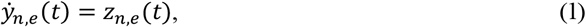

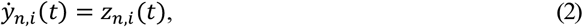

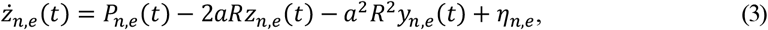

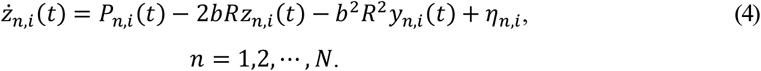

Here, *y*_*n,e*_, *y*_*n,i*_, *Z*_*n,e*_, and *Z*_*n,i*_ are excitatory and inhibitory PSPs, and excitatory and inhibitory post-synaptic current (PSC), respectively in the *n*^th^ brain region out of N brain regions depending on parcellation granularity. The subscripts *e* and *i* of the variables indicate excitatory and inhibitory populations, respectively. Parameters *a* and *b* are the reciprocal of the time constants of the PSP kernel for the excitatory and inhibitory populations. A random uniform distribution was used for independent noise *η*, and *R* scaled the spectral power distribution of the PSP signals.

The models of different brain regions were coupled through the excitatory populations, where the empirical structural connectome was employed to calculate the coupling strengths and delays forming a network backbone of the whole-brain model. The intra- and inter-region coupling is included in the terms *p*_*n,e*_ and *p*_*n,i*_ serving as inputs to the excitatory and inhibitory populations in region *n*, respectively, and had the following form:

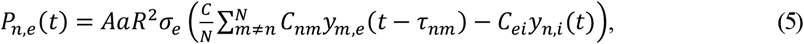

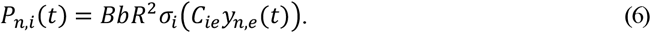

Parameters *A* and *B* are the maximal amplitudes of the excitatory and inhibitory PSP kernels. The inter-regional coupling strength and its delay from region *m* to region *n* (*C*_*nm*_ and *τ*_*nm*_) can be estimated using the empirical structural connectome

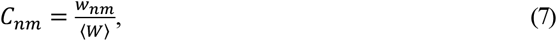

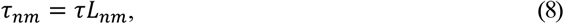

where *w*_*nm*_ and *L*_*nm*_ are the number and the average path-length of streamlines, respectively, between region *m* and region *n*, where the former was normalized by the averaged number of streamlines of the structural connectome ‹*W›*. Parameters of the global coupling *C* (arbitrary unit) and global delay *τ* (s/m) were to scale couplings and delays throughout the whole-brain network. The coupling weights *C*_*ie*_ and *C*_*ei*_ are balancing the interactions from excitatory to inhibitory populations and vice versa. The inter- and intra-region coupling involved an averaged firing density calculated by the following sigmoid functions:

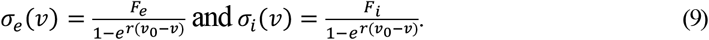

Here, *r* is a slope of the sigmoid function, *ν*_*0*_ is a half of the maximal membrane potentials. *F*_*e*_ and *F*_*i*_ are the maximal firing densities of the excitatory and inhibitory populations. The simulated excitatory PSP signals were applied to the neurovascular coupling and the hemodynamic function as described by the Balloon-Windkessel mode^l47,48^ that converted the simulated electrical neural activity to simulated BOLD signals. More details of the whole-brain model and their parameter values can be found elsewhere^22,39^,49.

### Implementation of simulation

The whole-brain model (1)-(9) was simulated by a custom-made program written in C++ with integration step of 2 ms during 720 s, where the first 57 s were discarded as a transient. The remaining 663 s (the same as the length of empirical rs-fMRI) were used for analysis. The simulation was carried out on the high-performance computing cluster^50^. The global coupling and global delay were varied as free model parameters on a dense grid of 64 values of global couplings [0, 63] and 43 values of global delays [0, 0.42] leading to 2752 model runs for each subject and parcellation.

### Network-based behavioral model fitting

Simulated BOLD signals were used to calculate the simulated FC by the pairwise Pearson’s correlation between the simulated BOLD signals of the brain regions. For the *behavioral network-based model fitting*, the graph-theoretical network properties (modularity and efficiency)^13^ of the simulated FC matrices were obtained for each model parameter point. To calculate the network properties, we took absolute values of the edges in the simulated FC matrices as for the empirical FC and delineated a parameter landscape (64-by-43 grid of model parameter points) using the network properties of the 2752 simulated FC. In the end, each subject had two network landscapes: modularity and efficiency for every parcellation. With this, group-level analyses and statistical tests were performed for statistical mapping on the landscapes across subjects. Under each condition (network property and atlas), parameter points were considered with sufficiently high inter-subject variability of the network properties. We therefore qualified the parameter points, where the standard deviations across subjects exceeded the third quartile of it over all parameter points (> 75%), which can positively contribute to investigation of the inter-individual differences and resistance against noise. Consequently, we can search for the optimal model parameters corresponding to the strongest correlation between the network properties and clinical scores (severity of the disease and disease duration) and the largest effect size of the group difference between healthy controls and PD patients. To see how the optimal parameter points are stable across different groups of sampled patients, we performed stratified 3-fold cross-validation for the behavioral network-based model fitting (200 iterations). Subsequently, Pearson’s correlation coefficients between disease severity and the considered network properties were calculated in training and testing steps.

### Statistical analysis

Effect sizes of group difference between healthy subjects and patients were calculated by the Rosenthal formula^51^ that used a *z*-statistic utilized to compute *p*-value of the non-parametric Wilcoxon rank-sum two-tail test (*n*=111). The statistics of the Pearson’s correlation coefficient between network properties and clinical or demographic variables were calculated for testing the hypothesis that there is no linear relationship (null hypothesis) between observations (*n*=60).

The Benjamini-Hochberg procedure^52^ was applied for controlling the false discovery rate (FDR) of a family of hypothesis tests from the edge-wise statistics that performed multiple times using the Pearson’s correlation (corrected *p*-values). The random-field theory^53^ for multiple tests was applied to statistical landscapes, and significant areas were thresholded (Z > 3.82 as corrected *p* < 0.05). The Kolmogorov-Smirnov test^54^ was used for normality of differences of the efficiency in healthy controls (*n*=51), and one-sample two-tail *t*-test was applied for testing the null hypothesis (no difference from zero) of the distribution. Statistical tests with *p* < 0.05 were considered as confirming the significance of results. The Benjamini-Hochberg FDR procedure was employed in Mass Univariate ERP Toolbox^55^. All statistical tests were performed in MATLAB (R2020b; MathWorks).

### Reporting summary

Further information on research design is available in the Portfolio Reporting Summary linked to this article.

## Data availability

The clinical data used in this study are not immediately available for public sharing because the given informed consent of the patients did not include public sharing. The simulated data that support the findings of this study are available from the corresponding author upon a reasonable request.

### Code availability

The brain connectivity toolbox is available here (https://sites.google.com/site/bctnet/). The containerized pipeline is publicly available (https://jugit.fz-juelich.de/inm7/public/vbc-mri-pipeline). Mass Univariate ERP Toolbox is a public software (https://openwetware.org/wiki/Mass_Univariate_ERP_Toolbox). The convolution-based neural mass model is also available from the first author upon a reasonable request.

## Acknowledgements

Primary contact of UKD-PD team is Prof. Dr. Julian Caspers. The authors gratefully acknowledge the computing time granted through JARA on the supercomputer JURECA at Forschungszentrum Jülich. This work was supported by the Portfolio Theme Supercomputing and Modeling for the Human Brain by the Helmholtz association, the Human Brain Project and the European Union’s Horizon 2020 Research and Innovation Programme under the Grant Agreements 785907 (HBP SGA2), 945539 (HBP SGA3) and 826421 (VirtualBrainCloud). Open access publication was funded by the Deutsche Forschungsgemeinschaft (DFG, German Research Foundation) - 491111487. The funders had no role in study design, data collection and analysis, decision to publish or preparation of the manuscript.

## Author contribution statements

K.J. contributed to conceptualization, data curation, formal analysis, investigation, methodology, resources, software, validation, visualization, writing - original draft, and writing - review & editing. S.B.E. contributed to conceptualization, funding acquisition, resources, project administration, and writing - review & editing. J.C. contributed to data curation and writing - review & editing. UKD-PD team contributed to data curation. O.V.P. contributed to conceptualization, funding acquisition, methodology, resources, project administration, software, supervision, validation, and writing - review & editing.

## Additional information

UKD-PD team

Julian Caspers^3^, Christian Mathys^3,4,5^, Martin Südmeyer^6,7^, Felix Hoffstaedter^1,2^, Christian J. Hartmann^6^, Christian Rubbert^3^, Alfons Schnitzler^6^, Bernd Turowski^3^

^4^Institute of Radiology and Neuroradiology, Evangelisches Krankenhaus Oldenburg, Universitätsmedizin Oldenburg, 26122 Oldenburg, Germany

^5^Research Center Neurosensory Science, Carl von Ossietzky Universität Oldenburg, 26129 Oldenburg, Germany

^6^Department of Neurology and Institute of Clinical Neuroscience and Medical Psychology, Medical Faculty and University Hospital Düsseldorf, Heinrich Heine University Düsseldorf, 40225 Düsseldorf, Germany

^7^Department of Neurology, Ernst-von-Bergmann Klinikum, 14467 Potsdam, Germany

## References

1 Kalia, L. V. & Lang, A. E. Parkinson’s disease. Lancet 386, 896–912, doi:10.1016/S0140-6736(14)61393-3 (2015).

2 DeLong, M. R. & Wichmann, T. Circuits and circuit disorders of the basal ganglia. Arch Neurol 64, 20–24, doi:10.1001/archneur.64.1.20 (2007).

3 Holden, S. K., Finseth, T., Sillau, S. H. & Berman, B. D. Progression of MDS-UPDRS Scores Over Five Years in De Novo Parkinson Disease from the Parkinson’s Progression Markers Initiative Cohort. Mov Disord Clin Pract 5, 47–53, doi:10.1002/mdc3.12553 (2018).

4 Gupta, H. V., Lyons, K. E., Wachter, N. & Pahwa, R. Long Term Response to Levodopa in Parkinson’s Disease. J Parkinsons Dis 9, 525–529, doi:10.3233/JPD-191633 (2019).

5 Santos-Garcia, D. et al. Response to levodopa in Parkinson’s disease over time. A 4-year follow-up study. Parkinsonism Relat Disord 116, 105852, doi:10.1016/j.parkreldis.2023.105852 (2023).

6 Ruppert, M. C. et al. Network degeneration in Parkinson’s disease: multimodal imaging of nigro-striato-cortical dysfunction. Brain 143, 944–959, doi:10.1093/brain/awaa019 (2020).

7 Steidel, K. et al. Longitudinal trimodal imaging of midbrain-associated network degeneration in Parkinson’s disease. NPJ Parkinsons Dis 8, 79, doi:10.1038/s41531-022-00341-8 (2022).

8 Espay, A. J. et al. Levodopa-induced dyskinesia in Parkinson disease: Current and evolving concepts. Ann Neurol 84, 797–811, doi:10.1002/ana.25364 (2018).

9 Manza, P., Amandola, M., Tatineni, V., Li, C. R. & Leung, H. C. Response inhibition in Parkinson’s disease: a meta-analysis of dopaminergic medication and disease duration effects. NPJ Parkinsons Dis 3, 23, doi:10.1038/s41531-017-0024-2 (2017).

10 Sporns, O., Tononi, G. & Kotter, R. The human connectome: A structural description of the human brain. PLoS computational biology 1, e42, doi:10.1371/journal.pcbi.0010042 (2005).

11 Bullmore, E. & Sporns, O. Complex brain networks: graph theoretical analysis of structural and functional systems. Nature reviews. Neuroscience 10, 186–198, doi:10.1038/nrn2575 (2009).

12 Bassett, D. S., Zurn, P. & Gold, J. I. On the nature and use of models in network neuroscience. Nature reviews. Neuroscience 19, 566–578, doi:10.1038/s41583-018-0038-8 (2018).

13 Rubinov, M. & Sporns, O. Complex network measures of brain connectivity: uses and interpretations. NeuroImage 52, 1059–1069, doi:10.1016/j.neuroimage.2009.10.003 (2010).

14 Sporns, O. Contributions and challenges for network models in cognitive neuroscience. Nat Neurosci 17, 652–660, doi:10.1038/nn.3690 (2014).

15 Griffa, A., Baumann, P. S., Thiran, J. P. & Hagmann, P. Structural connectomics in brain diseases. NeuroImage 80, 515–526, doi:10.1016/j.neuroimage.2013.04.056 (2013).

16 Crossley, N. A. et al. The hubs of the human connectome are generally implicated in the anatomy of brain disorders. Brain 137, 2382–2395, doi:10.1093/brain/awu132 (2014).

17 Fornito, A., Zalesky, A. & Breakspear, M. The connectomics of brain disorders. Nature reviews. Neuroscience 16, 159–172, doi:10.1038/nrn3901 (2015).

18 Medaglia, J. D., Lynall, M. E. & Bassett, D. S. Cognitive network neuroscience. J Cogn Neurosci 27, 1471–1491, doi:10.1162/jocn_a_00810 (2015).

19 Olde Dubbelink, K. T. et al. Disrupted brain network topology in Parkinson’s disease: a longitudinal magnetoencephalography study. Brain 137, 197–207, doi:10.1093/brain/awt316 (2014).

20 Zuo, C. et al. Global Alterations of Whole Brain Structural Connectome in Parkinson’s Disease: A Meta-analysis. Neuropsychol Rev, doi:10.1007/s11065-022-09559-y (2022).

21 Plaschke, R. N. et al. On the integrity of functional brain networks in schizophrenia, Parkinson’s disease, and advanced age: Evidence from connectivity-based single-subject classification. Hum Brain Mapp 38, 5845–5858, doi:10.1002/hbm.23763 (2017).

22 Jung, K. et al. Whole-brain dynamical modelling for classification of Parkinson’s disease. Brain Commun 5, fcac331, doi:10.1093/braincomms/fcac331 (2023).

23 Rubbert, C. et al. Machine-learning identifies Parkinson’s disease patients based on resting-state between-network functional connectivity. Br J Radiol 92, 20180886,doi:10.1259/bjr.20180886 (2019).

24 Owen, J. P. et al. The structural connectome of the human brain in agenesis of the corpus callosum. NeuroImage 70, 340–355, doi:DOI 10.1016/j.neuroimage.2012.12.031 (2013).

25 Goetz, C. G. et al. Movement Disorder Society-sponsored revision of the Unified Parkinson’s Disease Rating Scale (MDS-UPDRS): scale presentation and clinimetric testing results. Mov Disord 23, 2129–2170, doi:10.1002/mds.22340 (2008).

26 Caminiti, R. et al. Diameter, length, speed, and conduction delay of callosal axons in macaque monkeys and humans: comparing data from histology and magnetic resonance imaging diffusion tractography. J Neurosci 33, 14501–14511, doi:10.1523/JNEUROSCI.0761-13.2013 (2013).

27 Popovych, O. V., Manos, T., Hoffstaedter, F. & Eickhoff, S. B. What Can Computational Models Contribute to Neuroimaging Data Analytics? Front Syst Neurosci 12, 68,doi:10.3389/fnsys.2018.00068 (2019).

28 Gotts, S. J. et al. Two distinct forms of functional lateralization in the human brain. Proc Natl Acad Sci U S A 110, E3435–3444, doi:10.1073/pnas.1302581110 (2013).

29 Luo, C. et al. Decreased Resting-State Interhemispheric Functional Connectivity in Parkinson’s Disease. Biomed Res Int 2015, 692684,doi:10.1155/2015/692684 (2015).

30 Friston, K. J. et al. Statistical parametric maps in functional imaging: A general linear approach. Human Brain Mapping 2, 189–210, doi:10.1002/hbm.460020402 (1994).

31 Thompson, P. M. et al. Mapping cortical change in Alzheimer’s disease, brain development, and schizophrenia. NeuroImage 23 Suppl 1, S2–18, doi:10.1016/j.neuroimage.2004.07.071 (2004).

32 Hawrylycz, M. J. et al. An anatomically comprehensive atlas of the adult human brain transcriptome. Nature 489, 391–399, doi:10.1038/nature11405 (2012).

33 Andellini, M., Cannata, V., Gazzellini, S., Bernardi, B. & Napolitano, A. Test-retest reliability of graph metrics of resting state MRI functional brain networks: A review. J Neurosci Methods 253, 183–192, doi:10.1016/j.jneumeth.2015.05.020 (2015).

34 Domhof, J. W. M., Eickhoff, S. B. & Popovych, O. V. Reliability and subject specificity of personalized wholebrain dynamical models. NeuroImage 257, 119321, doi:10.1016/j.neuroimage.2022.119321 (2022).

35 Zuo, X. N. et al. Toward reliable characterization of functional homogeneity in the human brain: preprocessing, scan duration, imaging resolution and computational space. NeuroImage 65, 374–386, doi:10.1016/j.neuroimage.2012.10.017 (2013).

36 Owen, J. P. et al. Test-retest reliability of computational network measurements derived from the structural connectome of the human brain. Brain Connect 3, 160–176, doi:10.1089/brain.2012.0121 (2013).

37 Zhao, T. et al. Test-retest reliability of white matter structural brain networks: a multiband diffusion MRI study. Front Hum Neurosci 9, 59, doi:10.3389/fnhum.2015.00059 (2015).

38 Nakuci, J. et al. Within-subject reproducibility varies in multi-modal, longitudinal brain networks. Sci Rep 13, 6699, doi:10.1038/s41598-023-33441-3 (2023).

39 Jansen, B. H. & Rit, V. G. Electroencephalogram and visual evoked potential generation in a mathematical model of coupled cortical columns. Biol Cybern 73, 357–366, doi:10.1007/BF00199471 (1995).

40 Sip, V. et al. Characterization of regional differences in resting-state fMRI with a data-driven network model of brain dynamics. Sci Adv 9, eabq7547, doi:10.1126/sciadv.abq7547 (2023).

41 Tournier, J. D. et al. MRtrix3: A fast, flexible and open software framework for medical image processing and visualisation. NeuroImage 202, 116137, doi:10.1016/j.neuroimage.2019.116137 (2019).

42 Tournier, J. D., Calamante, F. & Connelly, A. Robust determination of the fibre orientation distribution in diffusion MRI: non-negativity constrained super-resolved spherical deconvolution. NeuroImage 35, 1459–1472, doi:10.1016/j.neuroimage.2007.02.016 (2007).

43 Schaefer, A. et al. Local-Global Parcellation of the Human Cerebral Cortex from Intrinsic Functional Connectivity MRI. Cereb Cortex 28, 3095–3114, doi:10.1093/cercor/bhx179 (2018).

44 Desikan, R. S. et al. An automated labeling system for subdividing the human cerebral cortex on MRI scans into gyral based regions of interest. NeuroImage 31, 968–980, doi:10.1016/j.neuroimage.2006.01.021 (2006).

45 Lopes da Silva, F. H., Hoeks, A., Smits, H. & Zetterberg, L. H. Model of brain rhythmic activity. The alpha-rhythm of the thalamus. Kybernetik 15, 27–37, doi:10.1007/BF00270757 (1974).

46 Moran, R., Pinotsis, D. A. & Friston, K. Neural masses and fields in dynamic causal modeling. Front Comput Neurosci 7, 57, doi:10.3389/fncom.2013.00057 (2013).

47 Buxton, R. B., Wong, E. C. & Frank, L. R. Dynamics of blood flow and oxygenation changes during brain activation: the balloon model. Magn Reson Med 39, 855–864, doi:10.1002/mrm.1910390602 (1998).

48 Friston, K. J., Mechelli, A., Turner, R. & Price, C. J. Nonlinear responses in fMRI: the Balloon model, Volterra kernels, and other hemodynamics. NeuroImage 12, 466–477, doi:10.1006/nimg.2000.0630 (2000).

49 Havlicek, M. et al. Physiologically informed dynamic causal modeling of fMRI data. NeuroImage 122, 355–372, doi:10.1016/j.neuroimage.2015.07.078 (2015).

50 Jülich Supercomputing Centre. JURECA: Data Centric and Booster Modules implementing the Modular Supercomputing Architecture at Jülich Supercomputing Centre. Journal of large-scale research facilities JLSRF 7, doi:10.17815/jlsrf-7-182 (2021).

51 Rosenthal, R., Cooper, H. & Hedges, L. Parametric measures of effect size. The handbook of research synthesis 621, 231–244 (1994).

52 Benjamini, Y. & Hochberg, Y. Controlling the False Discovery Rate: A Practical and Powerful Approach to Multiple Testing. Journal of the Royal Statistical Society: Series B (Methodological) 57, 289–300, doi:10.1111/j.2517-6161.1995.tb02031.x (1995).

53 Ashburner, J., Friston, K. J. & Penny, W. Human brain function. 2edn, (2003).

54 Massey, F. J. The Kolmogorov-Smirnov Test for Goodness of Fit. Journal of the American Statistical Association 46, 68–78, doi:10.1080/01621459.1951.10500769 (1951).

55 Groppe, D. M., Urbach, T. P. & Kutas, M. Mass univariate analysis of event-related brain potentials/fields I: a critical tutorial review. Psychophysiology 48, 1711–1725, doi:10.1111/j.1469-8986.2011.01273.x (2011).

