## Supplementary material for "Simulated brain networks reflecting progression of Parkinson’s disease"

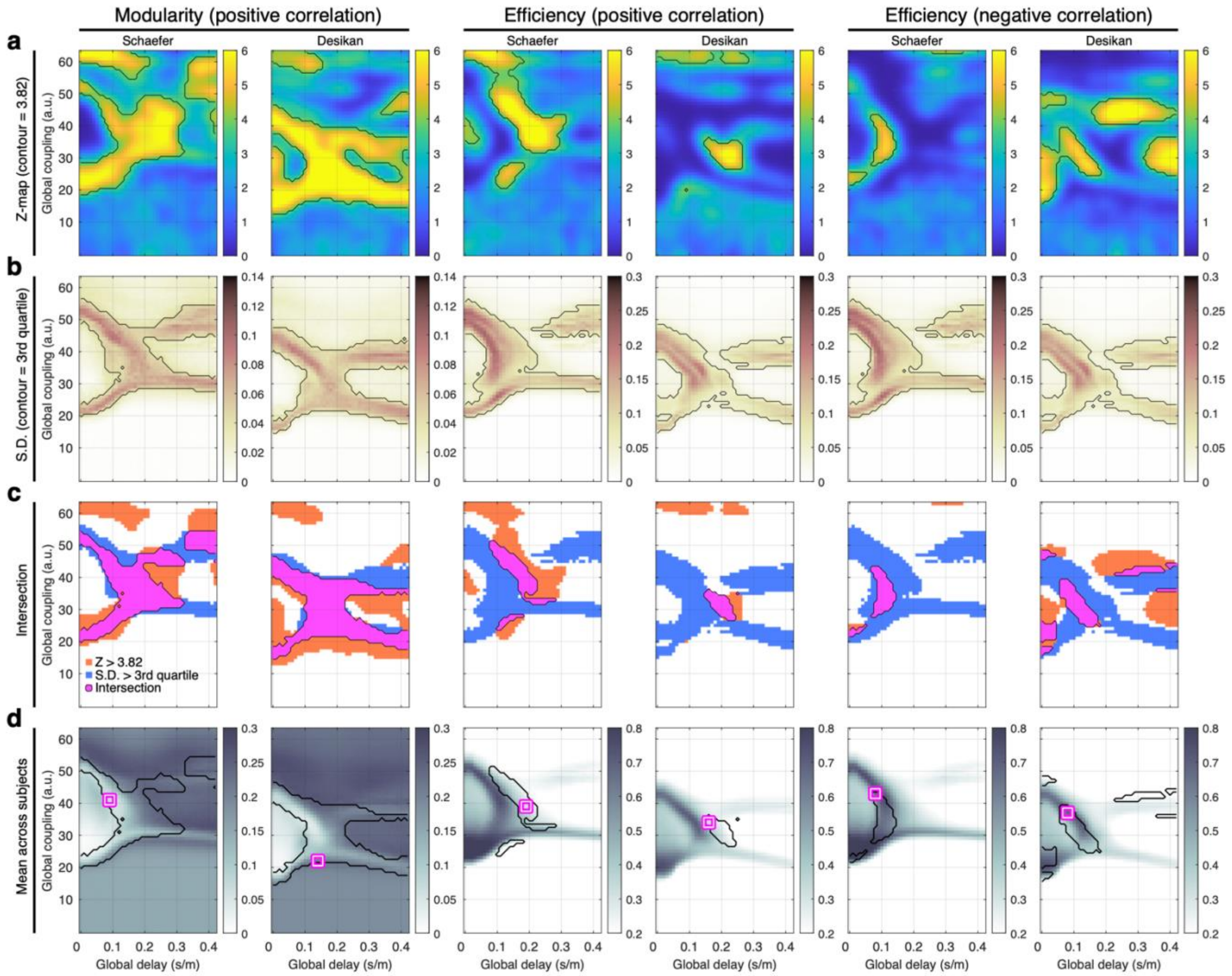


**Suppl. Fig. 1.** Parameter landscapes of statistical values for the computation conditions considered in the study. The network properties (modularity and efficiency) and brain parcellation schemes (Schaefer and Desikan-Killiany) are indicated in title of the plots. **(a)** Statistical maps for significant parameter areas of the Pearson's correlation coefficients between the network properties and severity of the disease (UPDRS III On: unified Parkinson's disease rating scale with medication). The black contour curves indicate the significant areas thresholded by the random-field theory for multiple tests (Z > 3.82 as corrected $p < 0.05$). **(b)** Statistical maps of the standard deviation (S.D.) of the network properties across subjects. The black contour curves indicate the areas of high inter-subject variance of the respected network properties (> third quartile). **(c)** Intersection in purple color of the significant areas from (a) and those of the high inter-subject variance from (b) of the respective network and parcellation conditions. **(d)** Landscapes of mean network properties (across the patients). The magenta-white squares indicate the optimal parameter points obtained and used in the main text and illustrated in Fig. 2c-e.

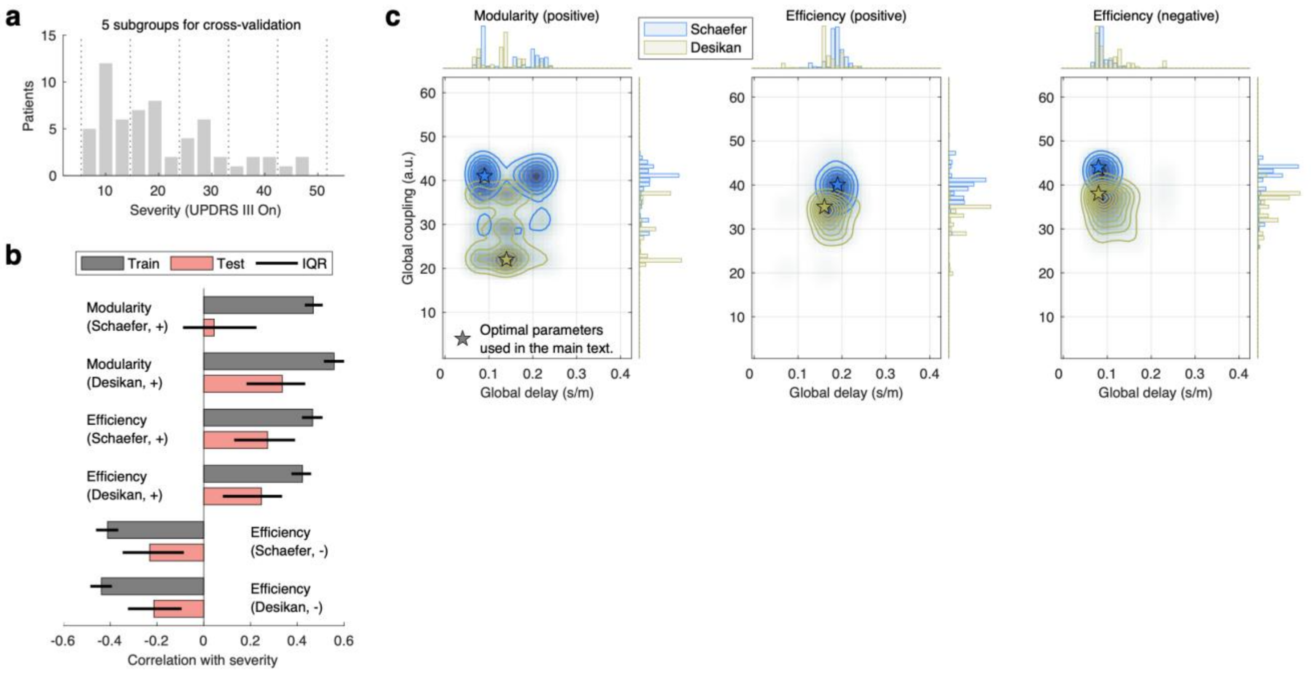


**Suppl. Fig. 2.** Stratified 3-fold cross-validation (CV) for the behavioral network-based model fitting with 200 iterations. **(a)** Stratified 5 subgroups for the CV keeping the distribution of the disease severity (UPDRS III On) in training and testing steps. **(b)** Pearson's correlation coefficients between disease severity and the considered network properties in the training and testing sets ($n$=200) for the considered brain parcellations and network properties of the simulated FC as indicated in the plot. The plus and minus signs correspond to the case of positive and negative correlations, respectively, between network properties and disease severity. **(c)** Distributions of the optimal model parameters derived by the cross-validated behavioral model fitting of the network properties of simulated FC to the disease severity for each parcellation and network property condition as indicated in the titles and legend. The stars indicate the optimal parameter points obtained and used in the main text and illustrated in Fig. 2c-e. The histograms on the top and right axes depict the distributions of the obtained optimal parameter values across CV.

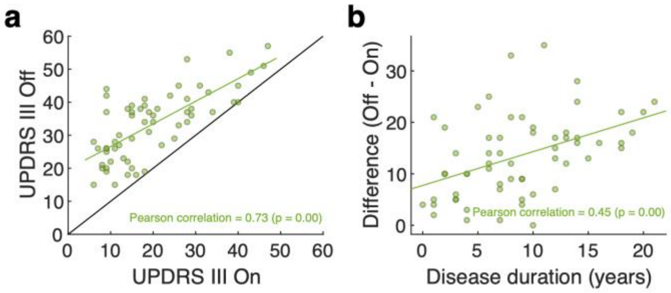


**Suppl. Fig. 3.** Scatter plots of **(a)** UPDRS III On versus UPDRS III Off and **(b)** disease duration versus difference of UPDRS III scores (Off − On). The empty circles in the plots correspond to individual subjects. The amount of correlation of the depicted relationships are indicated in these plots together with results of its statistical tests ($p$-values) of the Pearson's correlation.

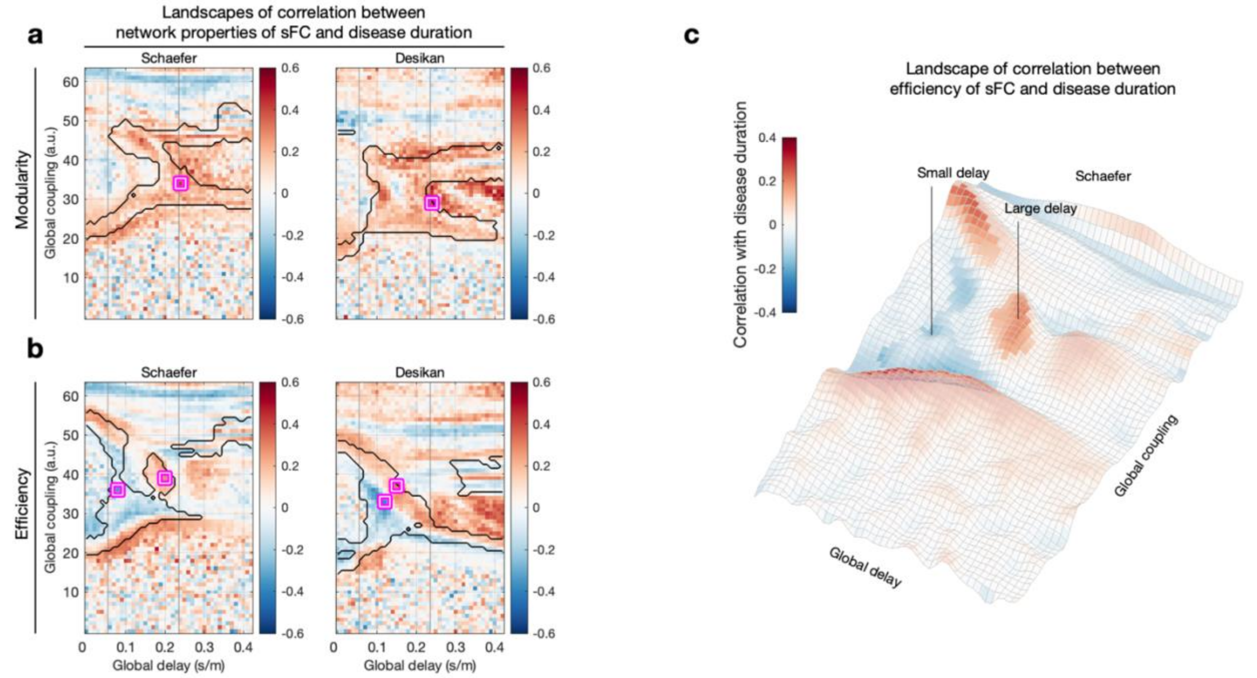


**Suppl. Fig. 4.** Parameter landscapes of the relationships between disease duration and the network properties of simulated FC used for the behavioral model validation. The network modularity (functional segregation) and network efficiency (functional integration) of the simulated FC were used to calculate **(a-b)** landscapes of Pearson's correlation (across PD patients) between simulated network properties and disease duration. The calculations were performed for the Schaefer and the Desikan-Killiany (Desikan) brain atlases indicated in the titles of plots together with the respective network properties. The vertical lines bound an approximate range of biologically feasible delays, the magenta-white squares indicate selected optimal parameter points of the correlation with disease duration in the parameter domain bounded by the black contour curves of intersection of significant areas thresholded by the random-field theory for multiple tests and areas of high inter-subject variance of the respected network properties (> third quartile). **(c)** Landscape of Pearson's correlation coefficients between simulated network efficiency and the disease duration in the Schaefer atlas. The vertical lines with 'small delay' and 'large delay' indicate selected optimal parameter points for negative and positive correlation, respectively.

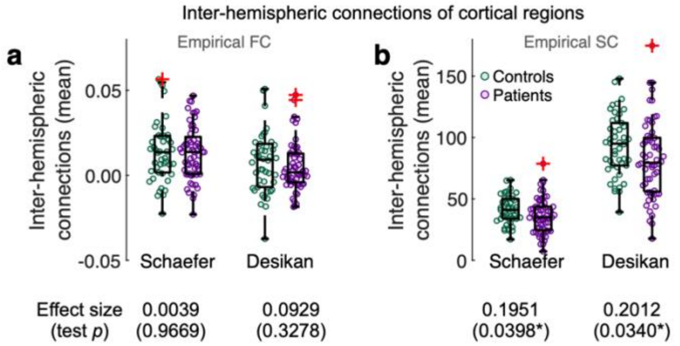


**Suppl. Fig. 5.** Inter-hemispheric connections of the empirical connectomes of two groups of PD patients and healthy controls. **(a)** Empirical functional connectivity (FC). **(b)** Empirical structural connectivity (SC). The brain parcellations are indicated in the plots, and the values under the plots are the effect sizes of the group difference (positive for HC > PD and negative for PD > HC) and their statistics (*p*-values of the Wilcoxon rank-sum two-tail test). The *p*-values with asterisks indicate significant results ($p < 0.05$). The red crosses away from the box plots are outliers.
